# SP1 and STAT3 functionally synergize to induce the RhoU small GTPase and a subclass of non-canonical WNT responsive genes correlating with poor prognosis in breast cancer

**DOI:** 10.1101/387951

**Authors:** Emanuele Monteleone, Valeria Orecchia, Paola Corrieri, Davide Schiavone, Lidia Avalle, Enrico Moiso, Aurora Savino, Ivan Molineris, Paolo Provero, Valeria Poli

## Abstract

Breast cancer is a complex disease in which heterogeneity makes clinical management very challenging. Although breast cancer subtypes classified according to specific molecular features are associated to better or worse prognosis, the identification of specific markers predicting disease outcome within the single subtypes still lags behind. Both the non-canonical WNT and the STAT3 pathways are often constitutively activated in breast tumors, and both can induce the small GTPase RhoU gene transcription. Here we show that RhoU transcription can be triggered by both canonical and non-canonical WNT ligands via the activation of JNK and the recruitment of the SP1 transcription factor to the RhoU promoter, identifying for the first time SP1 as a JNK-dependent mediator of WNT signaling. RhoU down-regulation by silencing or treatment with JNK, SP1 or STAT3 inhibitors lead to impaired cell migration in basal-like MDA-MB-231 cells, which display constitutive activation of both the non-canonical WNT and STAT3 pathways. These data suggest that STAT3 and SP1 can cooperate to induce high RhoU expression and enhance migration of breast cancer cells. *In vivo* binding of both factors characterizes a group of SP1/STAT3 responsive genes belonging to the non-canonical WNT and the IL-6/STAT3 pathways. High expression of this signature is significantly correlated with poor prognosis across all profiled patients. Thus, concomitant binding of both STAT3 and SP1 defines a subclass of genes contributing to breast cancer aggressiveness, suggesting the relevance of developing novel targeted therapies combining inhibitors of the STAT3 and WNT pathways or of their downstream mediators.

**Novelty and Impact:** The WNT and STAT3 pathways are often activated in breast tumors, but whether they can cooperate towards aggressiveness is not presently known. Here the authors show that WNT ligands can elicit the activation of the transcription factor SP1, which cooperates with STAT3 to induce a subset of non-canonical WNT and IL-6/STAT3 genes. Expression of this gene signature correlates with bad prognosis in breast cancer, suggesting coordinated interference with both TFs as a novel therapeutic option.

## INTRODUCTION

Breast cancer (BC) is a heterogeneous disease that is routinely classified according to the immunohistochemical assessment of Estrogen and Progesterone Receptors (ER, PR), human epidermal growth factor receptor 2 (HER2) and Ki67 [1, 2]. Alternatively, gene expression profiling according to the prediction analysis of microarray (PAM) 50 assay [3] can subdivide BC into intrinsic subtypes characterized by differential prognosis, i.e. luminal A, luminal B, HER2 positive, basal-like and normal-like. Gene signatures useful for informing therapeutic choices have been developed for early stage BC [3, 4], and the dependence on ER, PR or HER2 activity is currently exploited for the application of targeted therapies [1] which have considerably improved patients’ survival. However, the identification of specific markers responsible for worse prognosis across all cell types, potentially amenable to targeted therapies, is still needful.

WNT signaling is mediated by the WNT family of ligands, which controls fundamental processes in embryonic development and adult tissue homeostasis [5, 6]. According to the specific ligand involved, WNT signal transduction is broadly classified into canonical, β-catenin dependent, and non-canonical pathways, although the observation that specific co-receptors combinations in distinct cellular contexts can shape the pathway regardless of ligand identity challenges this rigid classification [7, 8]. Aberrant WNT signaling has been amply implicated in malignant transformation, and WNT ligands are overexpressed in a variety of cancers correlating with tumor progression [9, 10]. In particular, aberrant activation of the β-catenin-independent Planar Cell Polarity (PCP) pathway, which plays important roles in defining cell shape, adhesion and movement acting via the c-Jun N-terminal kinase (JNK) [6, 10], is involved in the progression of many solid tumours, including those of the breast [11, 12]. While transcriptional responses to the activation of beta-catenin-dependent WNT pathways are well characterized, those triggered by the PCP pathway are less well understood and have been mainly linked to the JNK-dependent activation of members of the AP1 family of transcription factors [13, 14],

The small GTPase RhoU was identified as a WNT1 target gene, mediating tumor transformation and promoting cell migration/invasion by inhibiting the formation of focal adhesions [15, 16]. Unlike most GTPases, RhoU displays high intrinsic guanine nucleotide exchange activity and is largely found in a GTP-loaded state [17, 18]. Accordingly, its regulation mostly occurs at the transcriptional level [19]. Despite WNT1 being considered a canonical ligand, WNT1-induced RhoU transcription is β-catenin-independent and requires JNK activity instead, suggesting the involvement of the non-canonical PCP pathway [19]. However, the molecular mechanisms leading to WNT-mediated RhoU transcriptional activation are currently unknown. RhoU gene expression can also be controlled by Signal Transducer and Activator of Transcription (STAT) 3 [19], a pleiotropic transcription factor activated downstream of many cytokines and growth factor receptors and playing a multitude of different roles in regulating cell proliferation/apoptosis, differentiation, migration and metabolism [20-22]. STAT3 is considered as an oncogene, since its aberrant constitutive activity is required for the survival and proliferation of a wide variety of primary tumors and tumor cell lines [20-24],

Here, we identify the SP1 transcription factor as an essential mediator of RhoU transcriptional activation downstream of the WNT/PCP pathway, and we unveil a functional cooperation between WNT/PCP/JNK, RhoU, SP1 and STAT3 to promote cell motility in basal-like human breast tumor cells. *In vivo* binding of both SP1 and STAT3 strikingly defines genes included in the non-canonical WNT and the IL-6 pathways. This novel signature is significantly correlated with low survival in breast cancer and is enriched in basal-like patients, suggesting that co-activation of the WNT-PCP and STAT3 pathways drives tumor aggressiveness.

## MATERIALS AND METHODS

### Cell lines and treatments

Stat3^+/+^ and Stat3^-/-^ immortalized MEF [25] and HEK-293 cells were cultured in DMEM (Dulbecco’s modified Eagle’s medium); MDA-MB-231 cells were purchased from ATCC and cultured in RPMI 1360 (Thermo Fisher Scientific, Waltham, Massachusetts,USA). Both media were supplemented with heat-inactivated 10% fetal bovine serum, 100 units/ml penicillin and 100 μg/ml streptomycin. WNT1/WNT4/WNT5a-expressing HEK-293 cells were obtained by stable transfection with vectors expressing the indicated WNT ligands. Cells were treated for 24 hours with the following inhibitors (Sigma-Aldrich, Saint Louis, MO, USA): 50 μM SP600125 (JNK inhibitor), 50 mM S3I-201 (STAT3 inhibitor) and 500 nM mithramycin A (SP1 inhibitor).

### Cell transfection and transduction, co-culture and luciferase assays

Mouse RhoU promoter fragments were generated by PCR using a genomic BAC vector as a template. The primers used are reported in Supporting Information Table 1. PCR products were cloned into the promoterless luciferase reporter vector pGL3 basic (Promega, WI, USA). Linker Scanning mutagenesis constructs, carrying 10 bp mutations starting from the indicated position (Supporting Information Fig. 1), were generated from the RhoU -756 promoter fragment. The primers used are shown in Supporting Information Table 2. The Super8X-TOPflash (TOP) and Super8X-FOPflash (FOP) constructs [6] were used to monitor WNT pathway activation.

Transient transfections, co-cultures with WNT-expressing cells (24 hours), luciferase and SEAP assays were performed essentially as previously described [19]. Briefly, transfections were performed with LIPOFECTAMINETM (Invitrogen, MA, USA) according to manufacturer’s instructions. Each construct was co-transfected at a 10:1 ratio with a vector expressing secreted embryonic alkaline phosphatase (SEAP) as an internal control for transfection efficiency.

MDA-MB-231 cells were stably infected with pLKO lentiviral vectors expressing two different shRNA sequences targeting RhoU (assays No 77507 and 48657, Thermo Fisher Scientific, Waltham, MA USA) or a scrambled sequence as a control, followed by puromycin selection (Sigma-Aldrich, Saint Louis, MO, USA, 1 μg/ml).

### Chromatin immunoprecipitatìon assays

ChIP experiments were performed as previously described [19] on three independent biological samples, using the following antibodies: anti-Sp1 sc-59X (Santa Cruz Biotechnology, Santa Cruz, CA); anti-histone H3 06-755 (Merck Millipore,Darmstadt, Germany); IgG (ChIP-IT Control Kit-Mouse; Active Motif, CA, USA). Non-immunoprecipitated chromatin was used as Total Input control (T.I.). Immunoprecipitated chromatin was analyzed by quantitative real-time PCR using Platinum SYBR Green PCR SuperMix-UDG with ROX (Life Technologies, Carlsbad, CA). Values were normalized to T.I. and to IgG signals. The β-globin promoter region was used as a negative control.

RhoU primers:

mRhoU-forward 5 ’-AGGGCAGGAGGAACTGGAGAGC -3’;
mRhoU-reverse, 5 ’-TACCCCTGGCCCCTGCTGTG-3 ’;
hRHOU-forward, 5 ’-ATAAAGGTTC ACGGC ATGCC-3 ’;
hRHOU-reverse, 5 ’-TAACTGCAGCTGATCGTGTG-3 ’;
β-globin primers:
β-glo-forward, 5 ’-CTCCCCCTCACTCTGTTCTG-3 ’;
β-glo-reverse, 5 ’-AGGAGGAGGGGA AGCTGAT A-3 ’.

### Wound healing migration assay

*In vitro* wound healing assays were performed essentially as described by Liang et al. [26], with some modifications. In brief, confluent cells in 60-mm dish were serum-starved 0/N, followed by mitomycin A treatment (10 μg/ml) for two hours. The monolayer was then scraped with a sterile pipette tip, washed with PBS and incubated in fresh medium containing 3% FBS. Phase-contrast images were taken at the indicated times (Axiovert 200M microscope, Carl Zeiss, Germany) (10X). Wound’s width was measured with the software ImageJ as the distance between the edges (National Institute of Health, Bethesda, Maryland, USA), and the migrating distance calculated as the difference between widths with respect to time 0, in 3 individual fields per experiment. As the edges were often irregular, the distance in each field was integrated from measures taken at three different position. Data are shown as mean±SEM of three independent experiments.

### Protein extracts and immunoblotting

Whole protein extractions and immunoblotting were performed as described [19] using the following antibodies: anti-Sp1 sc-59 (Santa Cruz Biotechnology, Santa Cruz, CA); mouse monoclonal anti-GFP and mouse monoclonal anti-vinculin (home-made production).

### RNA extraction, retrotranscription and quantitative real-time PCR (qRT-PCR)

These procedures were previously described [19], qRT-PCR reactions were performed with the ABI Prism 7300 realtime PCR System (Applied Biosystems, Carlsbad, CA) using Platinum Quantitative or SYBR Green PCR SuperMix-UDG with ROX (Life Technologies, Carlsbad, CA). Primers and probes used are reported in Supporting Information Table 3. The 18S rRNA pre-developed TaqMan assay (Applied Biosystems, Carlsbad, CA) was used as an internal control.

### Statistical analysis

Results were assessed for statistical significance by a standard two-tailed Student’s *t* test, using the GraphPad Prism 5 software (GraphPad Software Inc., San Diego, CA, USA). Statistically significant P values: P ≤ 0.05 (*), P ≤ 0.01 (**), P ≤ 0.001 (***).

### Bioinformatics Analysis

#### Total Binding Affinity (TBA)

TBA was computed as described in [27, 28] for all TFs included in the JASPAR core vertebrata database (version 4) with the promoter regions around the transcription start site (positions -756 to +235 bp, the same experimentally determined for the RhoU promoter) of all RefSeq genes (USCS genome browser version mm9).

#### Source data from database

mRNA normalized expression data and clinical feature were obtained by permission from the Molecular Taxonomy of Breast Cancer International Consortium (METABRIC, Syn1688369), downloaded at probe-level. Data from a total of 1981 breast invasive cancer tissue (BRCA) were obtained.

#### Chlp-Seq data analysis

ChIP-Seq data were downloaded from the UCSC browser [29]. Peaks were filtered according to the absolute distance (5 Kb) from the closest gene. Genes displaying binding sites for both TFs were selected.

*Gene set data.* Genes related to WNT and STAT3 signaling were obtained either from published manually curated gene sets [30-33], or from the “Molecular Signatures Database” (IL6, MsigDB C2) [34] (http://www.broadinstitute.org/gsea/msigdb/index.isp).

The SP1-S3 signature was defined as follows: genes defined by the ChIP-Seq data analysis were assessed for their enrichment with the above gene sets (Fisher exact test, p<0.05). A combined gene signature was obtained from the intersection between the two top-ranking signatures and the ChlP-Seq derived signature, obtaining 144 genes, listed in Supporting Information Table 4).

#### Survival analysis

Time-to-event analyses were performed using the Kaplan-Meier method (R, survival package). Patients were subdivided in two groups according to the median value of the SP1-S3 score, defined as the sum of SP1-S3 gene expression for each patient. Unstratified log-rank statistics were used to evaluate the effect of the change in SP1-S3 score on overall survival.

#### Univariate analysis

All univariate analyses were performed using the glm R function to assess the relationship between the SP1-S3 score and clinical features using the univariate Gaussian distribution. Logistic regression was applied for binary variables (https://www.R-proiect.org).

#### Plots

All plots were made using ggplot2 [35], survminer (https://CRAN.R-project.orq/packaqe=survminer) and R base graphics.

## RESULTS

### RhoU expression can be induced by both canonical and non-canonical WNT ligands

Our previous observation that the canonical WNT1 ligand triggers β-catenin-independent/JNK-dependent RhoU transcriptional induction prompted us to extend the analysis to other canonical or non-canonical WNT ligands. We thus measured RhoU mRNA in MEF cells stimulated with different WNT ligands, i.e. the canonical WNT1, the non-canonical WNT5a, and WNT4, which can activate both the canonical and the non-canonical pathway. Similar to WNT1, both WNT4 and WNT5a could trigger RhoU mRNA induction acting via the JNK-dependent PCP pathway, since the effect was abolished by the JNK inhibitor SP600125 (Figure 1A, B). In order to identify the transcription factors (TFs) involved, deletion fragments of the TATA-less RhoU promoter fused to a luciferase reporter gene were transiently transfected in MEF cells, followed by co-culturing with WNT-expressing or wild-type HEK-293 cells. All three WNT ligands could induce transcription of the RhoU promoter at similar levels, involving the same promoter region between positions -756 and -167 (Figure 1C). As expected, the β-catenin-dependent control TOPflash was solely responsive to the canonical WNT1 and WNT4 ligands. Further dissection identified a minimal region required for both basal and WNT-inducible promoter activity between positions -366 and -234 (Supporting Information Fig. 1A), conferring however weaker induction as compared to the sequence extending to position -756. Sequence analysis did not reveal binding sites for transcription factors know to be involved in WNT/PCP-mediated transcription such as those belonging to the AP1 family [19]. Therefore, in order to identify WNT-responsive elements we subjected the -756 construct Linker Scanning mutagenesis. Surprisingly, none of the mutations tested could affect promoter inducibility (Supporting Information Fig. 1B), suggesting that WNT-mediated RhoU induction does not depend on a single responsive element but requires the cooperation of disseminated sites within the responsive region.

**Figure 1.**
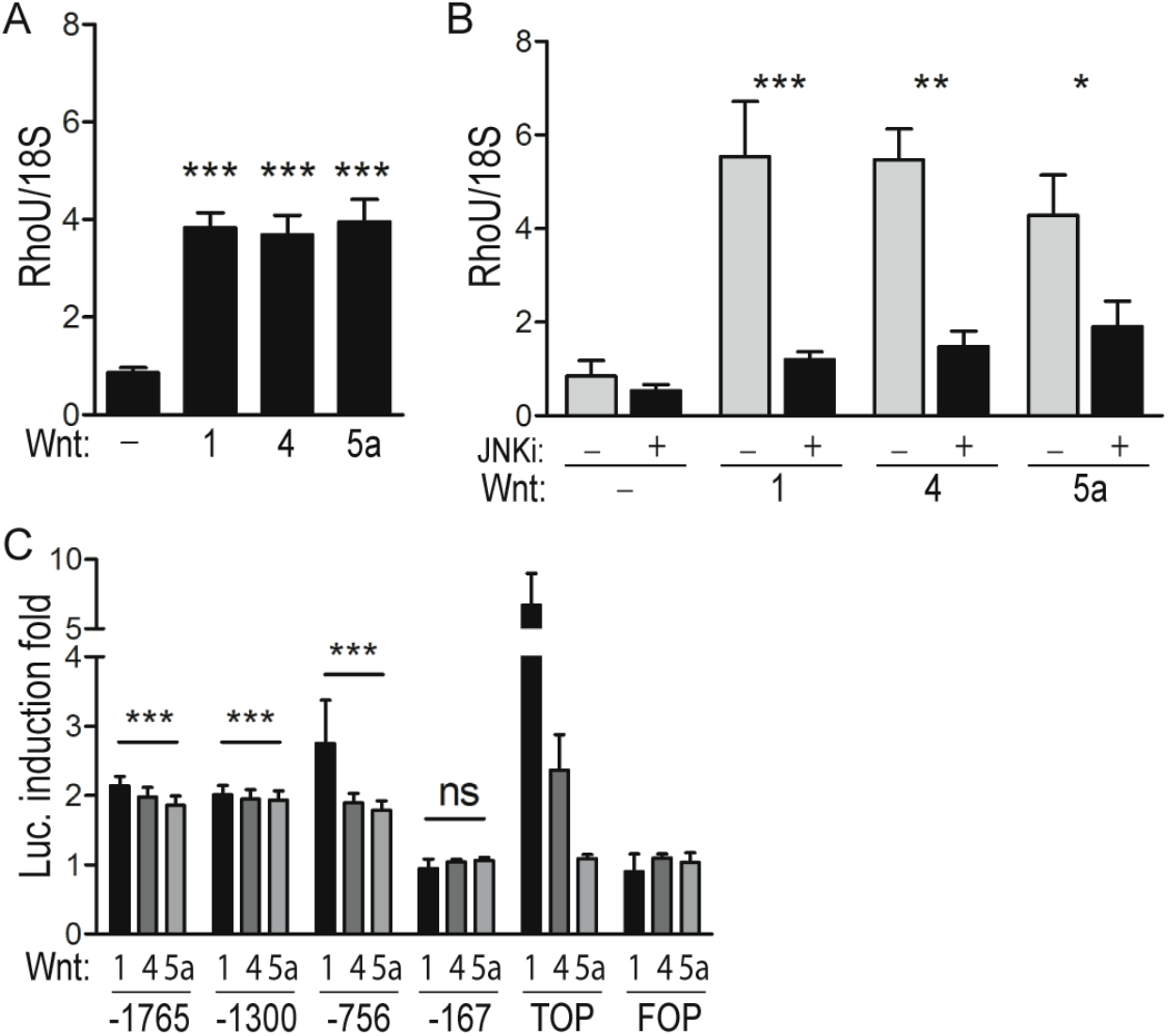
RhoU is a common WNT transcriptional target through the non-canonical JNKdependent pathway. (A, B) RhoU mRNA was measured by qRT-PCR in MEF cells co-cultured for 24 h with HEK-293 cells expressing or not the indicated WNTs (A), in the presence or absence of the JNK-inhibitor SP600125 (JNKi, B). Data are mean ± SEM of the relative expression levels, normalized to the 18S rRNA. N=3. (C) RhoU promoter activity in MEF cells transiently cotransfected with the indicated reporter constructs and a SEAP-expressing vector for normalization. TOPflash (TOP) and FOPflash (FOP) were used as positive and negative controls, respectively, for the β-catenin-dependent pathway. Transfected cells were stimulated or not with WNT ligands as described above. Data are mean ± SEM of the luciferase induction fold, relative to unstimulated condition. All WNT ligands significantly induced the activity of the -1765, -1300 and -756 RhoU promoter constructs (P<0.001), but not of the -167 unresponsive control (ns). *=P<0.05,**=P<0.01, ***=P<0.001; N=3.

### WNT-mediated induction of the RhoU promoter requires JNK-dependent SP1 recruitment

The failure to impair promoter inducibility with single mutations could be consistent with one or more TFs exhibiting diffused binding to the promoter due to high Total Binding Affinity (TBA), which considers the contribution of low-affinity binding sites to calculate the overall affinity of transcription factors for a chosen promoter region [27, 28, 36]. The TBA of the WNT-responsive RhoU promoter region for the TFs included in the JASPAR core vertebrata database was calculated (Supporting Information Table 5). Among high scoring TFs we focused our attention on SP1, known to play a major role in the transcription of GC-rich TATA-less promoters [37]. Although SP1 is mainly known as a basal activator of many housekeeping genes, it is also a target for JNK-ediated phosphorylation, which in turn can modulate its transcriptional and DNA binding activities at specific promoters [37-40]. We thus reasoned that WNT-activated JNK may mediate SP1 activation, enabling it to induce RhoU gene transcription via diffused binding to its promoter. SP1 involvement was confirmed by the observation that WNT-mediated RhoU mRNA induction was abolished by the SP1 inhibitor mithramycin A (Mit.A) (Figure 2A), which could also suppress the WNT1-mediated induction of the RhoU promoter but not of the TOPflash control (Figure 2B). Accordingly, ChIP analysis revealed strong *in vivo* binding of SP1 to the RhoU WNT-responsive promoter region following treatment with all three WNT ligands tested (Figure 2C, D). SP1 binding, undetectable in unstimulated cells, was abolished by both Mit.A treatment (Figure 2C) and JNK inhibition (Figure 2D). Thus, all tested WNT ligands can activate a non-canonical, JNK-mediated pathway that leads to SP1 recruitment to the RhoU promoter, driving its transcriptional activation. Accordingly, treatment with all three WNTs could increase the ratio of phosphorylated versus non-phosphorylated SP1 in a JNK-dependent manner, as shown by the relative increased intensity of the slower migrating band, corresponding to phosphorylated SP1 (Figure 2E, F). As expected Mit.A, which binds to GC-rich regions inhibiting SP1 DNA binding activity, did not affect WNT-mediated SP1 phosphorylation.

**Figure 2.**
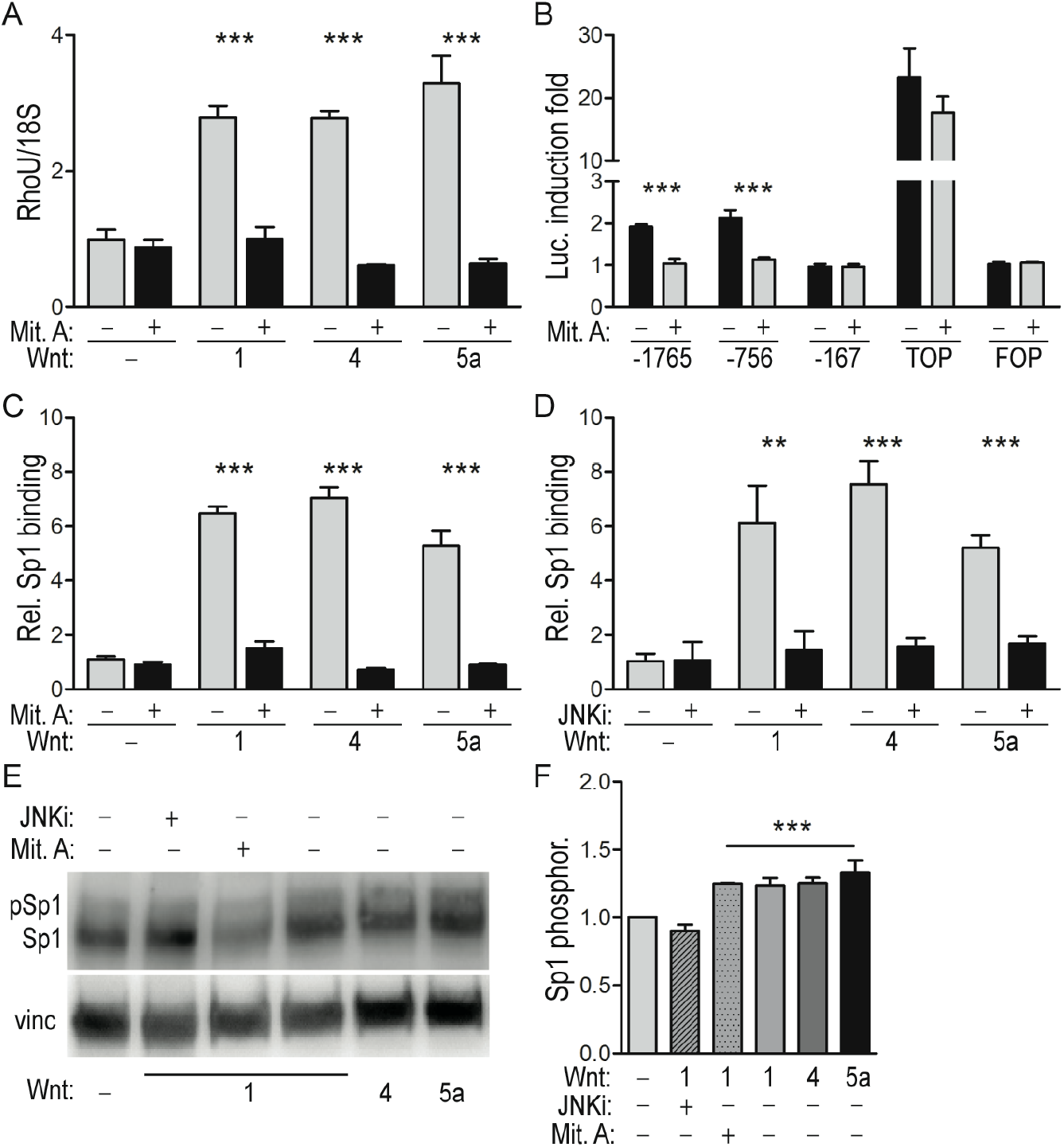
JNK-mediated SP1 recruitment is responsible for RhoU promoter induction by WNT ligands. (A) RhoU mRNA levels measured by qRT-PCR in MEF cells stimulated for 24 h with the indicated WNT ligands as described in the legend to Figure 1, in the presence or absence of the SP1 inhibitor Mithramycin A (Mit. A). Mean ± SEM of the relative expression levels normalized to the 18S rRNA; N=3. (B) The indicated RhoU promoter constructs were transfected in MEF cells as described in the legend to Fig. 1C, followed by WNT1 stimulation in the presence or absence of Mit. A. Data are mean ± SEM of the luciferase induction fold, relative to unstimulated condition. N=3. (C, D) SP1 binding to the RhoU promoter measured by ChIP assay in MEF cells stimulated for 24 h with the indicated WNTs, in the presence or absence of Mit. A (C) or of the JNK inhibitor JNKi (D). Immunoprecipitated chromatin was analyzed by qRT-PCR, and data (mean ± SEM) obtained upon normalization to total input (T.I.) and IgG. N=3. (E) Western blot showing WNT-dependent increase of phosphorylated SP1 (upper band). MEF cells were stimulated for 24 h with the indicated WNTs, in the presence or absence of Mit. A or JNKi, and analyzed with anti-Sp1 antibodies recognizing both the phosphorylated and unphosphorylated forms. Representative of one of four independent experiments. (F) Ratio between phosphorylated SP1 and total SP1 signals, expressed as fold induction relative to the unstimulated condition. Mean ± SEM of 4 independent experiments. N=4. **=P<0.01, ***=P<0.001.

### Migration of MDA-MB-231 cells downstream of constitutively active WNT5a requires both JNK and SP1 activities and RhoU expression

β-catenin-independent WNT signaling is often aberrantly activated in both tumor and stromal cells [11, 12, 41, 42], promoting JNK-dependent invasion [11, 42]. To assess the functional role of RhoU in this context we analyzed the highly aggressive basal-like breast cancer cell line MDA-MB-231, whose motility and invasivity have been shown to require the expression of the non-canonical WNT5a and WNT5b ligands [11, 42]. Indeed, as compared with the weakly motile MCF-7 cells, MDA-MB-231 cells express significantly higher levels of WNT5b and of the ROR1 co-receptor, implicated in the non canonical WNT pathway [43] (Figure 3A). Accordingly, RhoU expression was significantly higher in MDA-MB-231 cells, and fully dependent on both JNK and SP1 activities (Figure 3A,B). The motility of MDA-MB-231 cells in a wound healing assay could be significantly impaired both by shRNA-mediated silencing of RhoU (Figure 3C,D and Supporting Information Fig. 2A) and by treatment with the JNK or SP1 inhibitors (Figure 3E,F). Thus, SP1 activity is required to enhance cell migration downstream of the aberrant activation of the WNT/PCP/JNK pathway in basal-like cancer cells, at least partly via the induction of RhoU expression.

**Figure 3.**
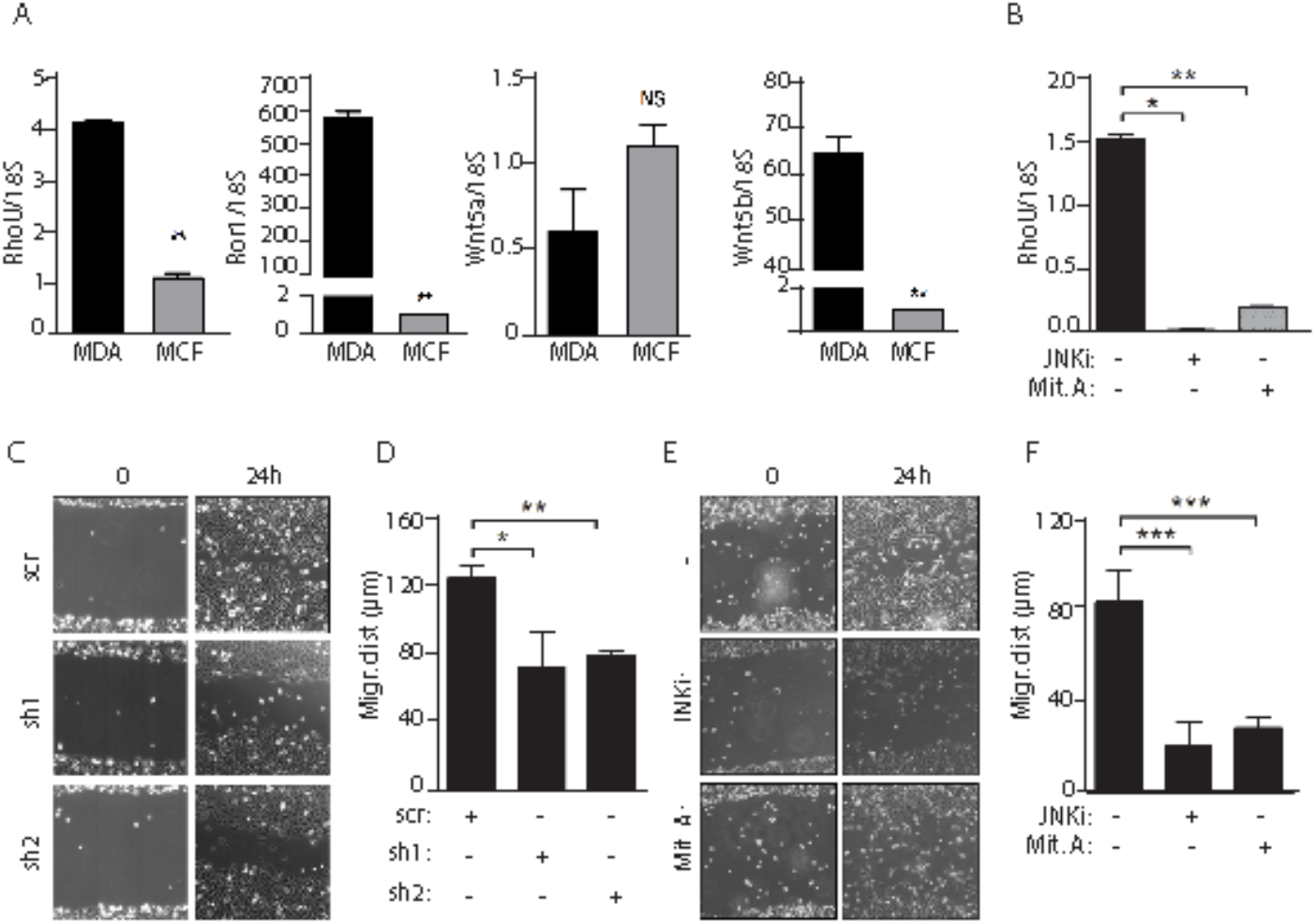
Both RhoU expression and JNK and SP1 activities are required for migration of MDA-MB-231 cells. (A) qRT-PCR analysis of RhoU, Ror1, Wnt5a and Wnt5b mRNAs in MDA-MB-231 and MCF-7 cells. (B) RhoU mRNA downregulation in MDA-MB-231 cells treated with the JNK or SP1 inhibitors (JNKi, Mit.A). Data are mean ± SEM of the relative expression levels, normalized to 18S rRNA. (C-F) Wound healing assays. MDA-MB-231 cells were infected with lentiviral vectors expressing two distinct RhoU shRNAs or a scrambled control (sh1, sh2, scr) (C, D), or treated with the JNK or SP1 inhibitors (JNKi, Mit.A) (E, F), and migration was assessed by *in vitro* wound healing assays. The phase contrast pictures (C, E) are representative of 3 independent experiments. Migrating distance (D, F) was calculated as described in the Materials and Methods, and shown as mean ± SEM. N=3. *=P<0.05, **=P<0.01, ***=P<0.001 between the indicated groups.

### RhoU is also involved in mediating cell migration downstream of STAT3 signaling

We recently identified RhoU as a STAT3 target gene downstream of gp130 cytokines, and reported a positive correlation between STAT3 activation and RhoU expression levels in a panel of human tumor cell lines, including MDA-MB-231 cells [19]. In order to investigate the role of RhoU in STAT3-induced migration, Stat3−/− MEFs, which express very low RhoU levels and display impaired migration compared to their wild-type counterparts [19, 44], were stably transfected with a doxycycline-inducible RhoU-GFP expression vector and treated with increasing doxycycline concentrations, obtaining dose-dependent RhoU induction (Supporting Information Fig. 2B). RhoU expression could significantly rescue the defective migration of Stat3−/− MEFs in a wound-healing assay (Figure 4A,B), demonstrating its involvement in STAT3-mediated migration. We then explored potential synergic effects between the WNT and STAT3 pathways in MDA-MB-231 cells. Treatment with the STAT3 inhibitor S3I considerably reduced the levels of RhoU mRNA (Figure 4C), leading to impaired cell migration at 24 hours, in synergy with the JNK inhibitor (P<0.01) (Figure 4D, E). These data are in line with the idea that RhoU indeed represents a common mediator of STAT3 and non canonical WNT/SP1 functions, under both physiological and pathological conditions.

### SP1 and STAT3 *in vivo* binding defines a subclass of genes belonging to non canonical WNT or IL-6 pathway

The TFs SP1 and STAT3, whose binding sites are often found close together in the regulatory regions of genes, are involved in several functions related to tumorigenesis including apoptosis, cell growth, angiogenesis, epithelial to mesenchymal transition (EMT) and invasion [45]. In order to investigate the STAT3 and SP1 regulatory network and to identify genes potentially regulated by both factors, we analyzed the ENCODE TFBS collapsed ChIP Sequencing track (wgEncodeRefTfbsClusteredV3) on H1-ESC, HeLa, MCF10A-Er-Src, K562, GM12878, HepG2 and HCT-116 cell lines, identifying ~3600 genes displaying ChIP-Sequencing peaks for both STAT3 and SP1 (see Materials and Methods section). While STAT3 binding could be detected on the RHOU promoter in MCF10A-Er-Src cells, the only mammary cell line available, which display active STAT3 [46], SP1 binding could not be detected in any of the cell lines analyzed. We reasoned that STAT3 and SP1 may coordinately mediate RHOU transcription in a cell type-specific manner, depending on the activation status of both factors. To expand our analysis, we took advantage of the “Integrative ChIP-Seq analysis of regulatory regions (ReMap 2018), which combines Public and ENCODE datasets, where we could detect *in vivo* SP1 binding to the RHOU promoter in HEK293T and A549 cells. The latter are lung adenocarcinoma cells displaying non-canonical WNT constitutive activity [47], and express high RhoU mRNA levels as detected by RNA sequencing. By ChIP-qPCR experiments we could demonstrate SP1 binding to the RhoU promoter in the MDA-MB-231 cells line, abolished in the presence of the SP1 inhibitor Mit.A (Fig. 5A). Enrichment analysis of the genes presenting both STAT3 and SP1 binding in published gene sets involving the activation of either the STAT3 or the WNT pathways revealed a positive and significant enrichment (p< 0.05, Fisher exact test) for eight out of seventeen gene sets (Fig. 5B). The ChIP-Seq traces for two representative genes are shown in Figure 5C. Of note, a high percentage of the peaks is located between positions −1000 and +1000 with respect to the TSS of the corresponding gene(s) (~74% for SP1, 57% for STAT3), compatibly with a regulatory role of the binding (data not shown). Importantly, the two top-ranking gene sets according to significance were linked to IL-6 (Broad Institute) and to non canonical WNT [31] signaling pathways, respectively (IL-6, ~37% overlap, 32 out of 87 genes; non-canonical WNT, ~23% overlap, 113 out of 489 genes).

**Figure 4.**
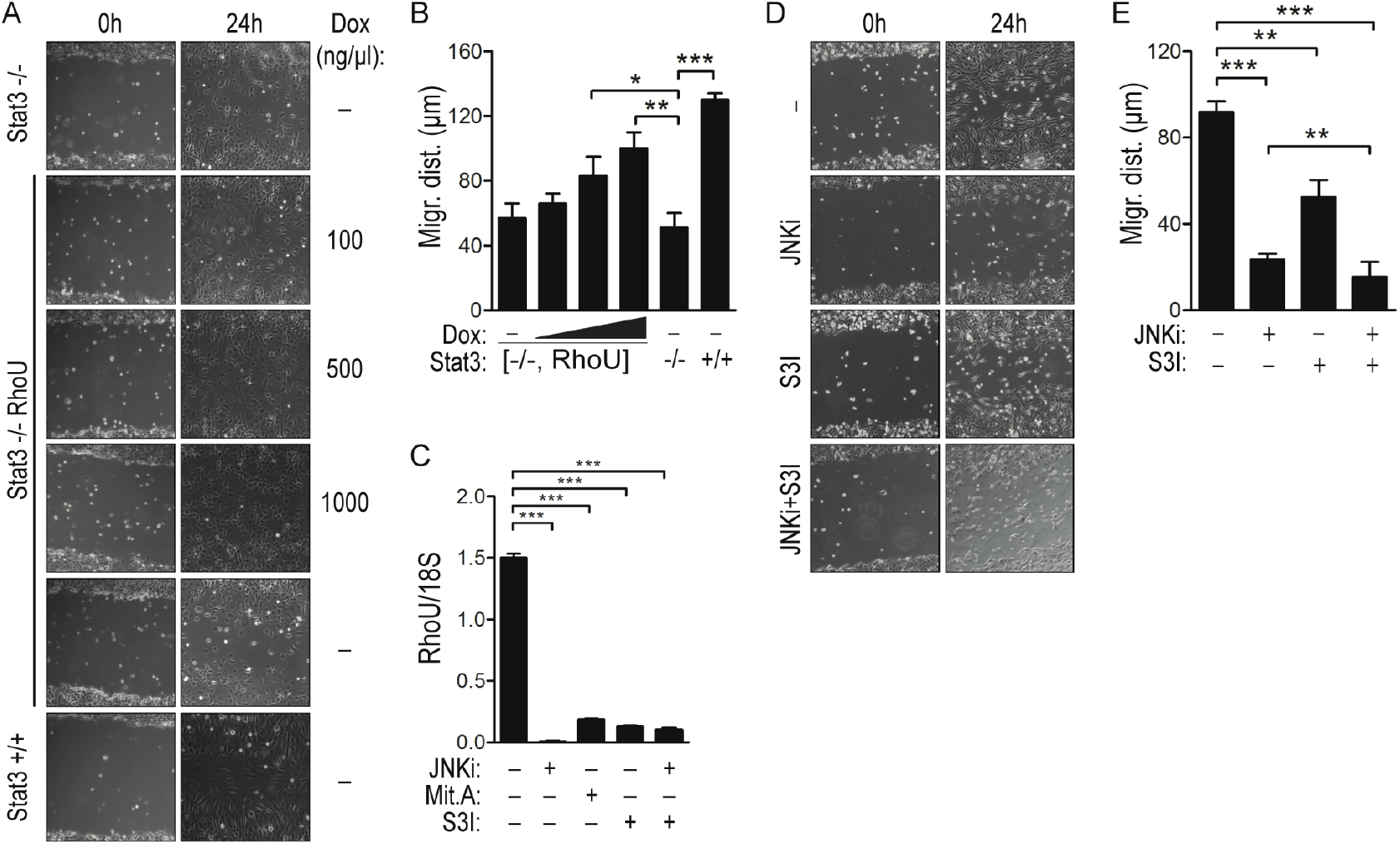
RhoU promotes cell migration also downstream of STAT3 in both MEF and tumor cells. (A, B, D, E) Wound healing assays were performed with the following cell types: wild type (Stat3^+^/+) or Stat3-deficient MEF cells treated or not with the indicated amounts of doxycycline (Dox) to induce increasing RhoU expression (Stat3−/­, Stat3−/­,RhoU) (A), and MDA-MB-231 cells treated with different combinations of the JNK and STAT3 inhibitors (JNKi, S3I) (D), and the migrating distance calculated as described in the legend to Figure 3 (B, E). (C) Downregulation of RhoU mRNA levels by the indicated inhibitors as measured by qRT-PCR in MDA-MB-231 cells. N=3. *=P<0.05, **=P<0.01, ***=P<0.001 between the indicated groups.

**Figure 5.**
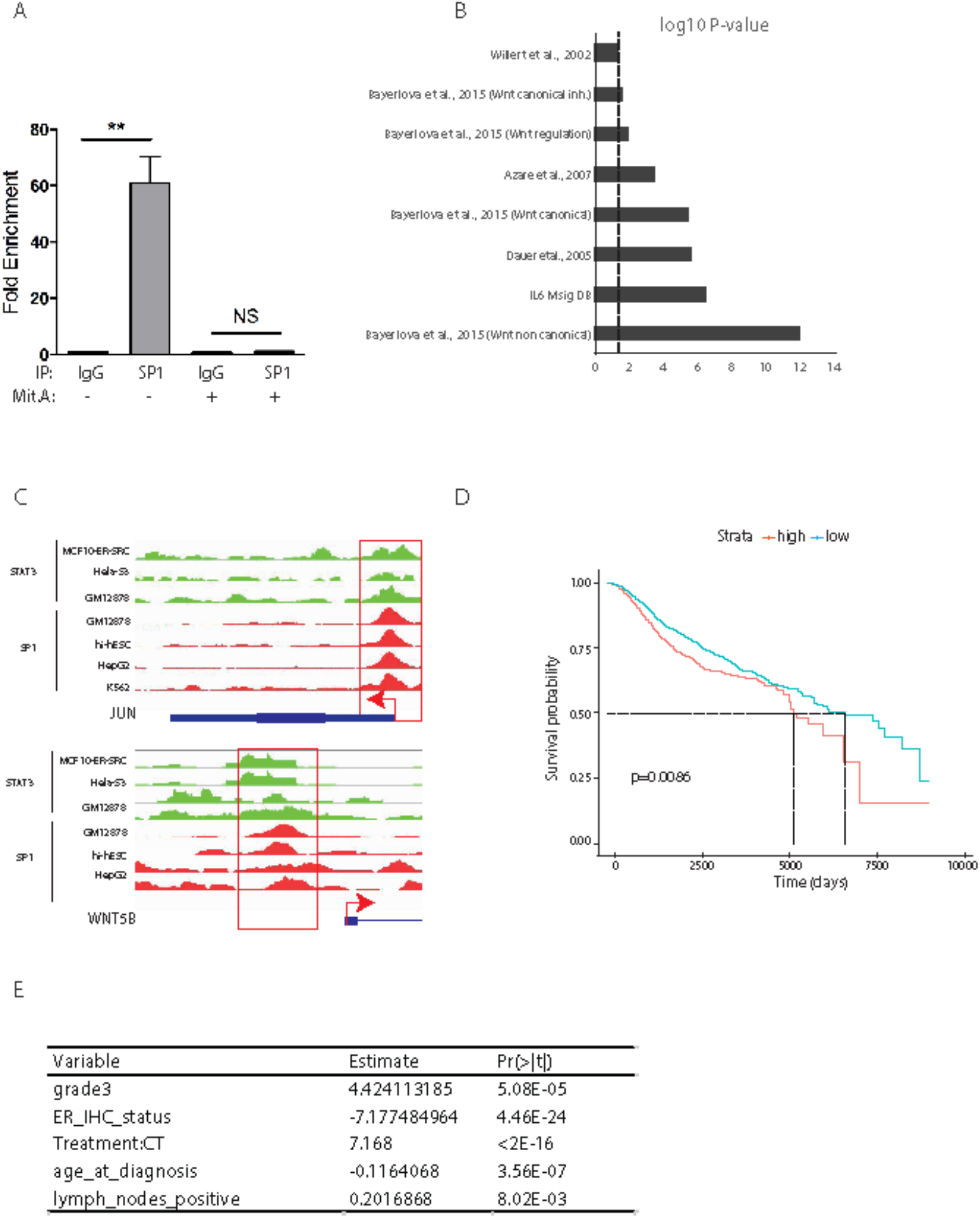
Concomitant activation of STAT3 and SP1 is associated with poor prognosis in breast cancer. (A) SP1 binding to the RhoU promoter measured by ChIP assay in MDA-MB-231 cells cultured in the presence or absence of Mit. A. Data are mean ± SEM of the fold enrichment obtained with anti-Sp1 antibodies versus IgGs in three independent experiments. (B) Bar plot showing the gene sets significantly enriched for STAT3 and SP1 bound genes. (C) IGV genome browser pictures showing binding of both SP1 and STAT3 in the indicated cell lines. Two representative genes of the two top-ranking gene sets in (B), WNT5B and JUN, are shown. (D) Kaplan-Meier plot of overall survival for breast tumor patients, as a function of time in days. Patient samples from the METABRIC database were subdivided according to low or high SP1-S3 score (median value). (E) Results of generalized linear models correlating grade, ER status, treatment, age at diagnosis and lymph node positivity to the SP1-STAT3 score. Coefficients (estimate) of the predictors and significance are indicated.

### High expression of SP1 and STAT3 dual targets is associated with low survival in breast cancer

We then assessed the functional relevance of combined STAT3 and SP1 activity in breast cancer. To this end, we examined the METABRIC breast cancer dataset [48] for the expression of a signature composed of the genes belonging to the IL-6 or the WNT non-canonical pathways characterized by *in vivo* binding of both STAT3 and SP1, as identified above, denominated the SP1-S3 signature (Supporting Information Table 4 and Table 6). All profiled tumors (n=1982) where classified according to the cumulative expression of the SP1-S3 signature (SP1-S3 score, high and low). This score was significantly higher in basal-like tumors compared to other subtypes, in keeping with the known frequent activation of both STAT3 and WNT pathways in this subset (Supporting Information Fig. 3 A). Patients with a high SP1-S3 score displayed significantly shorter overall survival as compared to patients scored as low (Fig 5D). To exclude that this correlation was due to the contribution of basal-like tumors, which can predict shorter survival when considered as a whole versus all other subtypes (not shown), we repeated the survival analysis excluding basal-like tumors. Even under these conditions, a high SP1-S3 score was still significantly predictive of poor prognosis, supporting the idea that SP1 and STAT3 can functionally cooperate to enhance tumor aggressiveness independently of the tumor subtype (Supporting Information Fig. 3B). A high SP1-S3 score was also significantly positively associated with tumor grade, higher degree of lymph node positivity and chemotherapy treatment, and negatively correlated with age at diagnosis and ER positivity (Fig 5E).

## DISCUSSION

Molecular classification of tumors based on gene signatures correlated with clinical history may guide prognosis, therapeutic choices and, finally, the development of novel specific therapies [4]. Both the WNT and the STAT3 pathways are often altered in solid tumors including breast cancer, correlating with enhanced tumor cells migration, invasion and metastasis [10, 49]. In an attempt to assess potential synergistic mechanisms between the two pathways, we focused on their common transcriptional target, the atypical small GTPase RhoU [19]. Our observation that RhoU induction via the JNK-dependent PCP pathway can be equally mediated by canonical and non-canonical WNT ligands suggests that this small GTPase may act as a common effector of both, expanding the scope for its potential relevance in mediating the pathogenic effects of disrupted WNT signaling.

Small GTPases belonging to the Rho family are well known to mediate downstream effects of the JNK/PCP pathway [50] that, although best characterized for its transcription-independent functions, is also known to lead to the transcriptional modulation of several genes, mainly via JNK-dependent activation of members of the AP1 family of transcription factors [13, 14]. The finding that RhoU is one of these transcriptional targets expands the repertoire of Rho GTPases downstream of the pathway. Moreover, our data identify SP1 as a novel transcriptional mediator of the JNK/PCP non canonical pathway, since its JNK-dependent recruitment to the RhoU promoter is essential for transcriptional induction downstream of both canonical and non-canonical WNT ligands. Although our data cannot discriminate between direct or indirect JNK-mediated SP1 activation, several pieces of evidence implicate this factor in inducible transcription upon both JNK and MAPK-mediated phosphorylation [38-40].

Our data establish a clear functional correlation between constitutive activation of the PCP pathway, JNK activity, SP1-mediated RhoU transcription and tumor cell motility. Indeed, RhoU downregulation dramatically impairs migration of the highly invasive basal-like MDA-MB-231 cells, which display constitutively high expression of the WNT5b ligand and of the ROR1 coreceptor, both implicated in non-canonical WNT pathways. Importantly, we show that also STAT3 converges on RhoU to support cell migration, since RhoU ectopic expression can partially rescue the defective migration of STAT3 null MEFs. Coordinated inhibition of both STAT3 and JNK cooperatively suppresses MDA-MB-231 cells migration, supporting the idea that the WNT/PCP/SP1 and STAT3 pathways can functionally synergize. Strikingly, the analysis of genes displaying *in vivo* binding for both SP1 and STAT3 detects a significant proportion of genes listed as non-canonical WNT or IL-6/STAT3 pathways, suggesting that the molecular mechanisms regulating RhoU transcription here identified may indeed extend to a subclass of WNT and IL-6/STAT3 responsive genes. The significant correlation of this gene signature with lower survival, high tumor grade, lymph node positivity, early age at diagnosis and chemotherapy treatment suggests that indeed the functional convergence of WNT/JNK/SP1 and STAT3 on downstream genes can confer aggressive features to breast tumors, independent of tumor subtype. Analysis of the genes of the signature reveals several interesting groups of genes, also exemplified by the results of the gene ontology analysis. Among the most enriched GO categories are “Chromatin Assembly”, “Chromatin Organization” and “DNA Metabolic Process”, including the genes encoding histone variants H2AX, essential for DNA repair and linked to chemoresistance in breast cancer [51] and H2AZ, whose high expression correlates with BC progression, lymph node metastasis and survival [52]. Other enriched categories are “Signal Transduction”, “Cell Communication” and “Response to Stimulus”, and include cytokine receptors involved in inflammation and immune response, integrins, important for migration and invasion, and several MAP Kinases. Indeed the category “MAPK Cascade” is also significantly enriched. MAP kinase genes showing dual SP1 and STAT3 binding are, for example, those encoding for MEK1, the proto-oncogenic MAP3K8 and p38, all well known players in Growth Factor and stress signaling linked to tumoral growth. Notably also JNK, the SP1 activator downstream of WNT, is a dual SP1-STAT3 target. Another significantly enriched category is “proteolysis”, and indeed it includes a striking number of proteasomal subunits, whose levels and activity are increased in a high percentage of primary breast cancers [53]. Among the “Regulation of Gene Expression, Epigenetic” and the “Transcription, DNA-dependent” categories are included many genes encoding subunits of the Mediator complex, often observed to contribute to the progression of several types of cancer [54], and several well known oncogenic transcription factors such as JUN and YAP. Finally, suggesting positive feed-forward mechanisms, SP1 itself as well as the WNT ligands WNT5b and WNT2b, the latter involved in both canonical and non-canonical signaling, are also dual SP1 and STAT3 targets.

Our data, shedding light on the molecular mechanisms regulating WNT/PCP-mediated transcriptional induction and pointing towards a cooperative co-regulation of many genes and gene families involved in tumor progression and aggressiveness by WNT/SP1 and STAT3, provide the rationale for efforts towards obtaining a coordinated inhibition of both pathways.

## ACKNOWLEDGMENTS

The authors wish to thank S.Oliviero and his group and G.Merlo (University of Torino) for helpful suggestions and generous gift of reagents, P.P.Pandolfi (University of Torino and Harvard) for helpful discussion, and S.Cabodi, P.Defilippi, F.Di Cunto, A.Camporeale and A.Camperi (University of Torino) for critically reading the manuscript.

This work was supported by the Italian Cancer Research Association (AIRC IG13009 and IG16930 to V.P.); the Italian Ministry of University and Research (MIUR PRIN to V.P. and P.P.); the Ateneo/Compagnia di San Paolo (to V.P. and P.P.), and the Truus and Gerrit van Riemsdijk Foundation, Liechtenstein, donation to V.P. E.M. was the recipient of a PhD fellowship from Fondazione CRT.

